# CTLA-4 blockade reverses the Foxp3+ T-regulatory-cell suppression of anti-tuberculosis T-cell effector responses

**DOI:** 10.1101/2020.05.11.089946

**Authors:** Lingyun Shao, Yan Gao, Xiaoyi Shao, Qingfang Ou, Shu Zhang, Qianqian Liu, Bingyan Zhang, Jing Wu, Qiaoling Ruan, Ling Shen, Xinhua Weng, Wenhong Zhang, Zheng W. Chen

## Abstract

**Backgrounds:** It has been well described that Foxp3+ T regulatory (Treg) cells suppress immune responses and that murine cytotoxic T lymphocyte-associated antigen 4 (CTLA-4) can control the function of Foxp3+Treg cells. However, it remains unknown about the role of CTLA-4 pathway in Treg suppression of T cell responses in tuberculosis (TB).

**Methods:** We assessed TB-driven changes in CTLA-4-expressing Foxp3+ Treg and conducted CTLA-4 blocking mechanistic studies ex vivo in 126 subjects with active TB, latent TB or uninfected statuses.

**Results:** Frequencies of CTLA-4-expressing Treg cells were increased in the circulation of pulmonary TB patients and in the pleural compartment of TB pleuritis. Six-month anti-TB treatment significantly reduced CTLA-4+ Treg subset. Notably, antibody blocking of CTLA-4 pathway (CTLA-4 blockade) reversed the ability of Treg cells to suppress anti-TB Th1 responses and abrogated the Treg-mediated suppression of TB antigen-stimulated proliferative response. The CTLA-4 blockade reversed the Treg suppression of the ability of T cells to restrict intracellular BCG and *M. tuberculosis* growth in macrophages.

**Interpretation:** The study uncovered previously-unreported observations implicating that the CTLA-4 blockade abrogates the capability of Treg cells to suppress anti-TB immune responses or immunity. Findings support the rationale for exploring the CTLA-4 blockade as potential host-directed therapy against TB.

**Fund:** This work is supported in part by the National Natural Science Foundation of China (30901277, 81671553, 81501359), and the Key Technologies Research and Development Program for Infectious Diseases of China (2017ZX10201302-004).

## Introduction

Tuberculosis (TB) remains a leading killer among infectious diseases despite that the World Health Organization (WHO) initiated the End TB Strategy in 2014. Clearly, efforts to control and eradicate TB are hampered by coinfection with the human immunodeficiency virus (HIV) and emergence of drug-resistant TB (1, 2). Doubtlessly, global control of TB will require better prevention and therapeutic strategies. Identification of host factors involving TB development or progression may help to discover adjunct host-directed therapies capable of boosting current treatment protocols(3).

Studies in animal models and humans suggest that TB-specific T-cell effector functions are impaired during active TB and that inhibitory factors may play a role in TB pathogenesis (4–7). In fact, it has been shown that Foxp3+ T regulatory cells (Treg) can suppress TB-specific T-cell responses *ex vivo* (8, 9). Foxp3+ Treg cells appear to exhibit the following 2 cardinal features: (i). Foxp3+ Treg cells constitutively express cytotoxic T lymphocyte antigen 4 (CTLA-4) (10–13); (ii). Foxp3 controls the expression of CTLA-4 in Treg cells(14–16).

CTLA-4 is a member of the B7/CD28 family and is a molecular antagonist of CD28. CTLA-4 and CD28 molecules mediate opposing functions in T-cell activation while binding to the same ligand B7 (CD80/86), but the CTLA-4 binding affinity is 500-2500 times higher than CD28. CD28 is expressed on all T cells, while surface CTLA-4 expression was limited to Foxp3+ Treg cells and some activated T cells. The CD28-B7 binding in conjunction with T-cell receptor (TCR) signals promotes T cell proliferation and IL-2 production. In contrast, the CTLA-4-B7 binding inhibits the transmission of co-stimulatory signals, thereby suppresses the proliferation and differentiation of antigen-specific T cells (12, 17, 18). It has been shown in CTLA-4 knockout mice that CTLA-4 is a key molecular target for controlling Treg-suppressive function in both physiological and pathological conditions including autoimmunity, allergy, and tumor immunity(13).

The CTLA-4 pathway is increasingly targeted as part of immune modulatory strategies to treat cancers, generically termed as immune checkpoint blockade for reversing an abnormal immune signaling. Targeting the CD28/CTLA-4 pathway using antibodies and fusion proteins is of considerable interest in the treatment of cancer and autoimmune diseases (17, 19, 20). While still in early stages, data from basic and clinical studies suggest that antibody blocking of CTLA-4 pathway (CTLA-4 blockade) can be beneficial for the treatment of chronic HIV, HBV, and HCV infections (21, 22). However, little is known about whether CTLA-4 plays a role in TB and the CTLA-4 blockade can potentially serve as host-directed therapy against TB. To gain new information, we assessed TB-driven changes in CTLA-4-expressing Foxp3+ Treg cells and conducted CTLA-4 blocking mechanistic studies ex vivo in 126 subjects with active TB, latent TB and uninfected statuses. We focused on our efforts to investigate whether the CTLA-4 blockade could reverse the Treg-mediated suppression of anti-TB Th1 responses and intracellular *M. tuberculosis* growth control.

## Materials and Methods

See the online supplement for an extended version of Methods.

### Study Participants

One hundred and twenty-six individuals with different tuberculosis infection status recruited in this study. Each participant was given written informed consent before blood testing. Clinical Information was recorded including age, gender, history of prior active TB, chest radiography, sputum acid-fast smear, sputum culture and other medical examinations. All individuals had a history of newborn Bacille Calmette-Guerin (BCG) vaccination.

Subjects were divided into three groups according to the statuses of tuberculosis infection. (1). Active tuberculosis group (ATB; n=75); (2) Latent tuberculosis infection group (LTBI; n=18); (3) Healthy uninfected control group (HC; n=33) (supplemental text). Demographic and clinical characteristics of study patients are shown in Table 1.

**Table 1.** Demographic and clinical characteristics of study subjects

|  | Total | ATB group |  |  | LTBI group | HC group | P value |
| --- | --- | --- | --- | --- | --- | --- | --- |
|  |  | Subtotal | PTB | TP |  |  |  |
| **Patient number** | 126 | 75 | 56 | 19 | 18 | 33 | -- |
| **Male/female (% of male)** | 68/58 (40.3) | 47/28 (62.7) | 34/22 (60.7) | 13/6 (68.4) | 7/11 | 14/19 | 0.811 |
| **Median Age, year (Range)** | 39.8 (18-85) | 42.6 (18-85) | 41.9 (18-80) | 44.8 (18-85) | 36.5 (22-65) | 35.1 (18-56) | 0.972 |
| **Previous TB history, n (%)** | 8 (6.3) | 8 (10.7) | 6 (10.7) | 2 (10.5) | 0 | 0 | <0.001 |
| **AFB smear positive, n (%)** | 58 (46.0) | 58 (77.3) | 56 (100.0) | 2 (10.5) | 0 | 0 | <0.001 |
| ***M.tb* Culture positive, n (%)** | 59 (46.8) | 59 (78.7) | 56 (100.0) | 3 (15.8) | 0 | 0 | <0.001 |
| **Anti-TB treatment, n (%)** | 51 (40.5) | 51 (68.0) | 42 (75.0) | 9 (47.4) | 0 | 0 | <0.001 |
| **BCG vaccination history, n (%)** | 101 (80.2) | 57 (76.0) | 42 (75.0) | 15 (81.3) | 14 (77.8) | 30 (90.9) | 0.072 |

ATB: active tuberculosis; PTB: pulmonary tuberculosis; TP: tuberculous pleurisy; LTBI: latent tuberculosis infection; HC: healthy control; AFB: acid fast bacilli; BCG: Bacille Calmette-Guerin

The study was approved by the Ethics Committee of Huashan Hospital, Fudan University, and written informed consent was obtained from all the participants.

### T-SPOT.TB assay

T-SPOT.TB assay was performed according to the manufacturer’s instructions as previously described(23–25) (supplemental text).

#### Isolation of peripheral blood mononuclear cells (PBMCs) and pleural fluid mononuclear cells (PFMCs)

Isolation of peripheral blood mononuclear cells (PBMCs) performed as previously described(26). (supplemental text)

#### Immunofluorescent staining and flow cytometric analysis

For cell surface staining, 100μL EDTA blood was treated with red blood cell (RBC) lysing buffer and washed twice with 5% fetal bovine serum (FBS)-phosphate-buffered saline (PBS) before staining. PBMCs stained with up to four Abs for at least 10 min at room temperature. After stained with cell-surface CD3/CD4/CD8/CD25, cells were permeabilized for 45 min at 4°C and stained another 45 min for CD152 (CTLA-4)-Biotinylated, then another 45 min for FoxP3, Streptavidin-PB.

Intracellular cytokine staining (ICS) performed as previously described (26). Briefly, 10^6^ cells plus mAbs CD28 (1μg/mL) and CD49d (1μg/mL) were incubated with PPD (25μg/mL) or media alone for 1 h at 37° C, 5% CO_2_ followed by an additional 5 h incubation in the presence of brefeldin A. After stained with cell-surface CD3/CD4/CD8, cells were permeabilized for 45 min at 4°C and stained another 45 min for IFN-γ-PE and IL-10-FITC at 4°C (supplemental text).

After staining, cells analyzed by a BD FACS Aria flow cytometer, at least 20 000-gated events were analyzed using FCS Express V3 Software.

### MACS sorting of Treg and Tresp

PBMCs samples using MACS CD4+ CD25+ CD127-/Low selection kits, according to manufacturer instructions. The unlabeled cell fraction of Tresp (CD4+CD25-), and the magnetically labeled cells Tregs (CD4+CD25+CD127^dim^) were got from PBMCs. Tregs and Tresp cells purities were routine > 95% (supplemental text).

### CFSE-labeled T lymphocyte proliferation inhibition assay

T cells were labeled with 0.5μM carboxyfluorescein diacetate succinimidyl ester (CFSE) just before stimulation. In the cells, esterases cleaved the acetyl group, leading to the fluorescent diacetylated CFSE. Cell division accompanied by CFSE dilution was analyzed by flow cytometry.

### Real-time PCR

Total RNA was isolated with TRIzol reagent (Life Technologies) by following the manufacturer’s instruction. For the reverse transcription reaction, TaqMan reverse transcription reagents (Life Technologies) used as described(27, 28). The relative quantities of mRNAs were determined by using the comparative Ct method and were normalized by using human glyceraldehydes-3-phosphate dehydrogenase (GAPDH) as an endogenous control (supplemental text).

### CTLA-4 blocking experiments

Tresp (CD4+CD25-) and Tregs (CD4+CD25+CD127^dim^) from PBMCs were cultured at a 1:2 Treg to Tresp ratio. Cell cultures were incubate with isotype control antibody or CD152(CTLA-4)-F(ab’)2 (10μg /ml) plus PPD (25μg /ml). After three days of incubation in RPMI medium supplemented with 10% (vol/vol) FBS, cells were stained with antibodies and analyzed by BD FACS Aria flow cytometer, and harvested with TRIZol for real-time PCR detection as described above. For the CFSE-labeled T lymphocyte proliferation inhibition assay, cells were incubated for 7 days and at 3^rd^ day plus IL-2, and at the 7^th^ day, cells were collected and stained with antibodies and analyzed by BD FACS Aria flow cytometer.

### *M. tuberculosis* and BCG culture *and in vitro* intracellular Mycobacterium growth inhibition assay (MGIA)

*M. tuberculosis* (*M.tb*) H37Rv and BCG were cultured at 37 °C in Middlebrook 7H9 medium containing 10% v/v OADC supplement and 0.05% w/v Tween 80 plus 100 mg/mL hygromycin B (supplemental text). The H37Rv-infected experiment was performed in the BSL-3 laboratory in Fudan university.

### Statistical analysis

The nonparametric quantitative variables across groups were compared by use of the Mann-Whitney test or paired *t* test. and qualitative variables by chi-square test. The relationship between clinical and demographic characteristics was assessed by ANOVA. Significance was inferred for *P*<0.05 with two-sided, and all analyses were done with Stata 7.0 software. And the graphs were drew by using GraphPad Prism 5.01 software.

## Results

### CTLA-4 expression in CD4+CD25+Foxp3+ Treg cells was increased both in the circulation of pulmonary TB patients and in the pleural compartment of TB pleurisy

As an initial effort for studies of CTLA-4 pathway and Treg cells function, we examined CTLA-4 expression in Foxp3+ Treg cells in TB. Little is known about CTLA-4 expression in Foxp3+ Treg cells during TB infection of humans, although TB-driven increases in Treg cells are well described (5, 29). Thus, CD4+CD25+Foxp3+ T (Treg) cells and CTLA-4 expression in Treg cells were comparatively investigated in 24 patients with active TB, 18 individuals with latent TB infection and 33 uninfected healthy controls using the flow cytometry-based analysis. ATB group exhibited significantly higher frequencies of circulating Foxp3+Treg cells than LTBI and HC groups, respectively (*P*=0.0119 and 0.0082, respectively) (Figure 1). Consistently, ATB group displayed significantly higher frequencies of the CTLA-4-expressing Foxp3+Treg subpopulation than LTBI or HC group (both *P*<0.0001), with median percentages of CTLA-4+Foxp3+ Treg subset being 22.85%, 6.99% and 9.54%, respectively (Figure 2).

**Figure 1.**
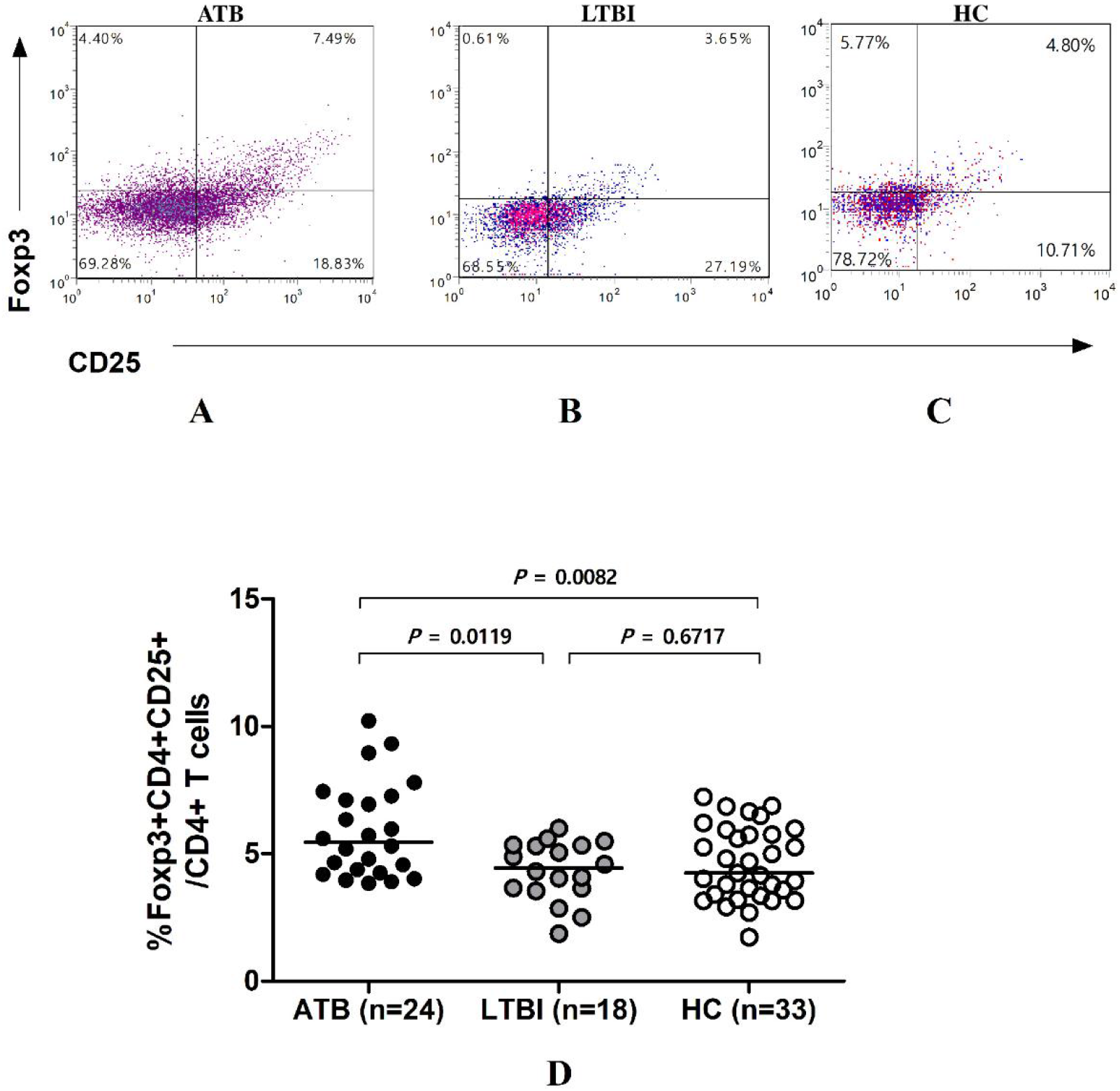
Frequencies of CD4+CD25+Foxp3+ Treg cells in subjects with different TB infection statuses. (A), (B) and (C) are the representative CD4-gated flow cytometry histograms of CD4+CD25+Foxp3+ Treg cells from an ATB patient, a LTBI individual and a HC. (D) The percentages of CD4+CD25+Foxp3+ Treg cells among CD4+ T cells. The horizontal lines represent the medians for each group. ATB: active TB (n=24), Twenty-four independent experiments were performed with comparable result; LTBI: latent TB infection (n=18). Eighteen independent experiments were performed with comparable result; HC: healthy controls (n=33). Thirty-three independent experiments were performed with comparable result. The nonparametric quantitative variables across groups were compared by use of the Mann-Whitney test. Significance was inferred for P<0.05 with two-sided.

**Figure 2.**
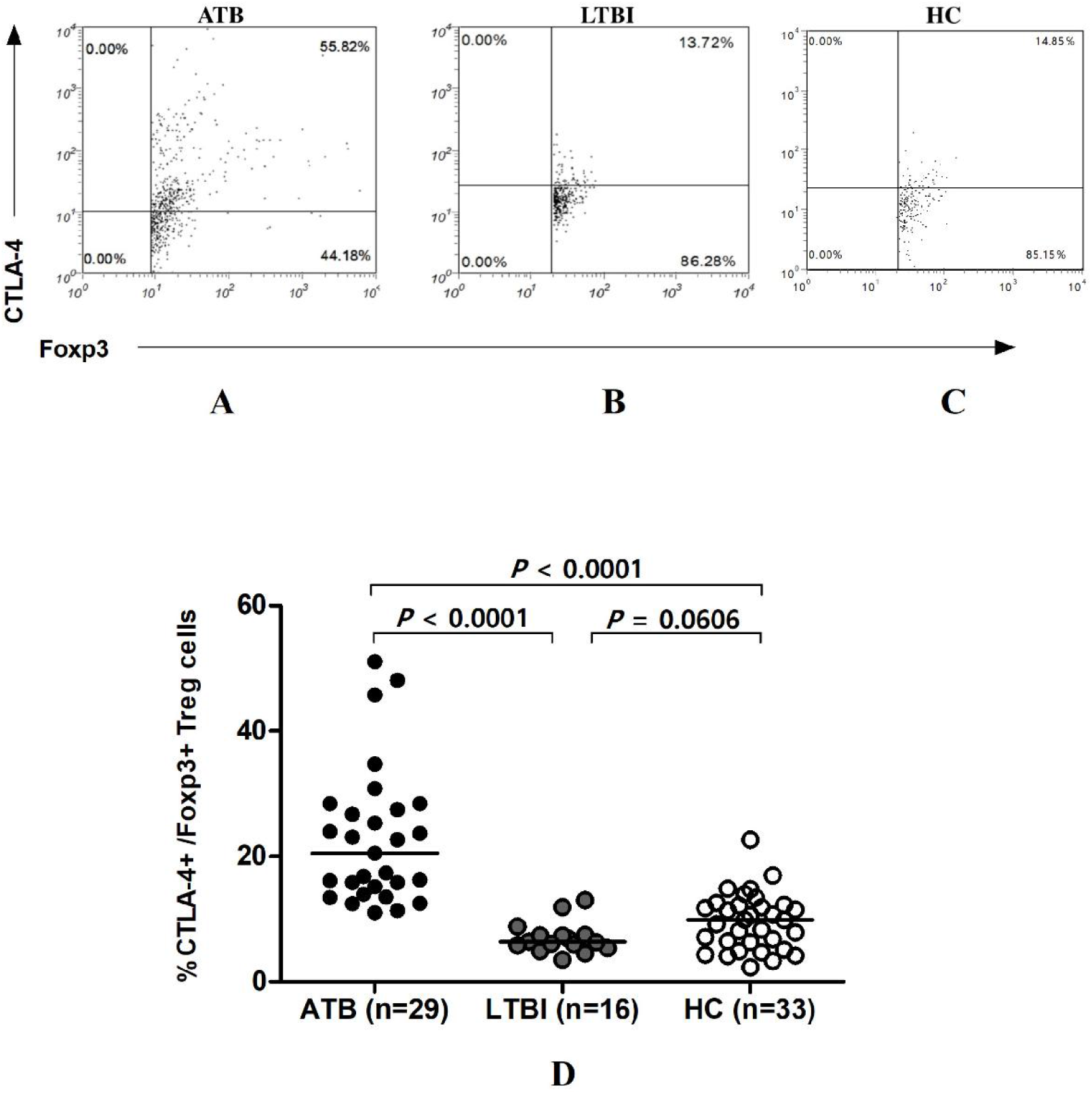
Frequencies of CTLA-4-expressing Foxp3+ Treg in subjects with different TB infection status. (A), (B) and (C) show the representative CD4CD25-gated flow histograms of CTLA-4+Foxp3 Treg cells from an ATB patient, a LTBI individual and HC. (D) shows the percentages of CTLA-4+Foxp3+ Treg cells. The horizontal lines represent the medians for each group. ATB: active TB (n=29), Twenty-nine independent experiments were performed with comparable result; LTBI: latent TB infection (n=16); HC: healthy controls (n=33). Thirty-three independent experiments were performed with comparable result. The nonparametric quantitative variables across groups were compared by use of the Mann-Whitney test. Significance was inferred for P<0.05 with two-sided.

In parallel, we recruited patients with TB pleurisy to examine whether CTLA-4 expression in Foxp3+Treg cells was up-regulated in the pleural infection site. Blood and pleural fluid were simultaneously collected from 13 TB pleurisy patients and comparatively assessed for frequencies of Foxp3+ Treg cells and CTLA-4+Foxp3+ Tregs. Interestingly, frequencies of both Foxp3+ Treg and CTLA-4+Foxp3+ Treg cells in pleural fluid mononuclear cells (PFMCS) were significantly higher than those in PBMCS from TB pleurisy patients (Figures 3A, 3B, *P*<0.01 for both cell subsets in blood-pleurisy comparisons)(5). Consistently, immunohistochemistry analysis showed that a number of Foxp3+ and CTLA-4+ lymphocytes were infiltrated in the pleural tissue collected from TB pleurisy patients (Figure S1).

**Figure 3.**
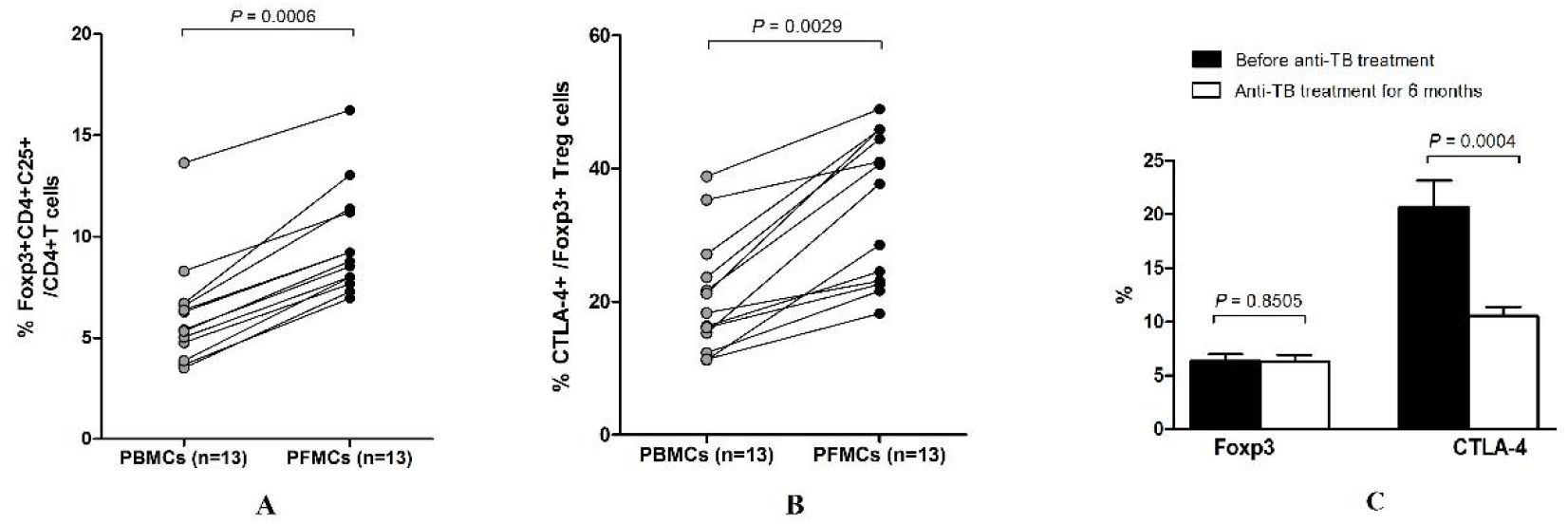
Frequencies of Foxp3+ Treg and CTLA-4+Foxp3+ Treg cells in PBMCs and PFMCs of patients with tuberculous pleurisy. (A) The percentages of CD4+CD25+Foxp3+ Treg cells among CD4 + T cell (n=13). (B) The percentages of CTLA-4+ Treg cells among CD4+CD25+Foxp3+ Treg cells (n=13); PBMCs: peripheral blood mononuclear cells; PFMCs: tuberculous pleural effusion mononuclear cells. (C) Changes in frequencies of Foxp3+Treg and CTLA-4+ Treg cells in PBMCS of patients with tuberculous pleurisy before and 6 months after anti-TB treatment (n=13). 13 patients with tuberculous pleurisy were enrolled for this assay and the independent experiments were performed with comparable result. Y axis shows the percentage of Foxp3+CD4+CD25+ Treg in CD4+ T cells or CTLA-4+ in FoxP3+CD4+CD25+ Treg cells. The nonparametric quantitative variables across groups were compared by use of the paired t test. Significance was inferred for P<0.05 with two-sided.

Together, these results suggest that CTLA-4-expressing Foxp3+ Treg cells are increased both in the circulation of pulmonary TB patients and in the pleural compartment of TB pleurisy.

### Anti-TB treatment significantly reduced TB-driven elevation of CTLA-4+Foxp3+ Treg cells

We then sought to determine if anti-TB treatment could alter TB-driven increases in CTLA-4+Foxp3+ Tregs. We followed up 16 patients with tuberculous pleurisy for immune studies before and 6 months after anti-TB treatment. CTLA-4+Foxp3+ Treg cells were significantly reduced from 20.69% to 10.55% in PBMCS at 6 months after anti-TB treatment (*P*=0.0004, Figure 3C), although there was no apparent change in frequencies of Treg cells. Such close relationship between CTLA-4 expression and TB infection well justified the need to investigate whether CTLA-4 pathway was involved in Treg-related immune suppression of anti-TB immune responses in the context of CTLA-4 blockade.

### The CTLA-4 blockade abrogated the ability of Foxp3+ Tregs to suppress PPD-stimulated IFN-γ T effector response, and such reversing effect did not coincide with reduction of IL-10

We then sought to explore whether CTLA-4 pathway plays a role in Treg suppression of anti-TB T-cell responses. It is noteworthy that there is a lack of a reliable conditional knock-out technique leading to CTLA-4 deficiency selectively in primary human Foxp3+Treg or T effector subsets. We therefore conducted CTLA-4 blocking mechanistic studies of T-cell subsets in TB patients. We asked the straight forward question as to whether antibody blocking of CTLA-4 pathway (CTLA-4 blockade) could block Foxp3+ Treg suppression of anti-TB T effector responses ex vivo in TB. This question has not been addressed in the field, and is highly practical or relevant to exploring the CTLA-4 blockade for potential immune intervention against TB.

Thus, PBMCS from 10 patients with active TB were employed to isolate CD4+CD25+CD127^-/dim^ T-cell subset (termed Treg) and CD4+CD25-T-cell subset (termed Tresp) using the standard MACS separation kit. These two cell subsets were cultured with PPD in different ratio in the presence or absence of CTLA-4 blockade. In the absence of CTLA-4 blockade, addition of purified Treg cells to the culture clearly suppressed the ability of CD4+CD25-Tresp cells to produce IFN-γ in response to PPD stimulation (Figure 4). In the presence of CTLA-4 blockade, however, such Treg-mediated suppression of Tresp production of IFN-γ was significantly reversed (Figure 4, *P*=0.0003), suggesting that CTLA-4 pathway involves Treg suppression of TB-specific T-cell responses.

**Figure 4.**
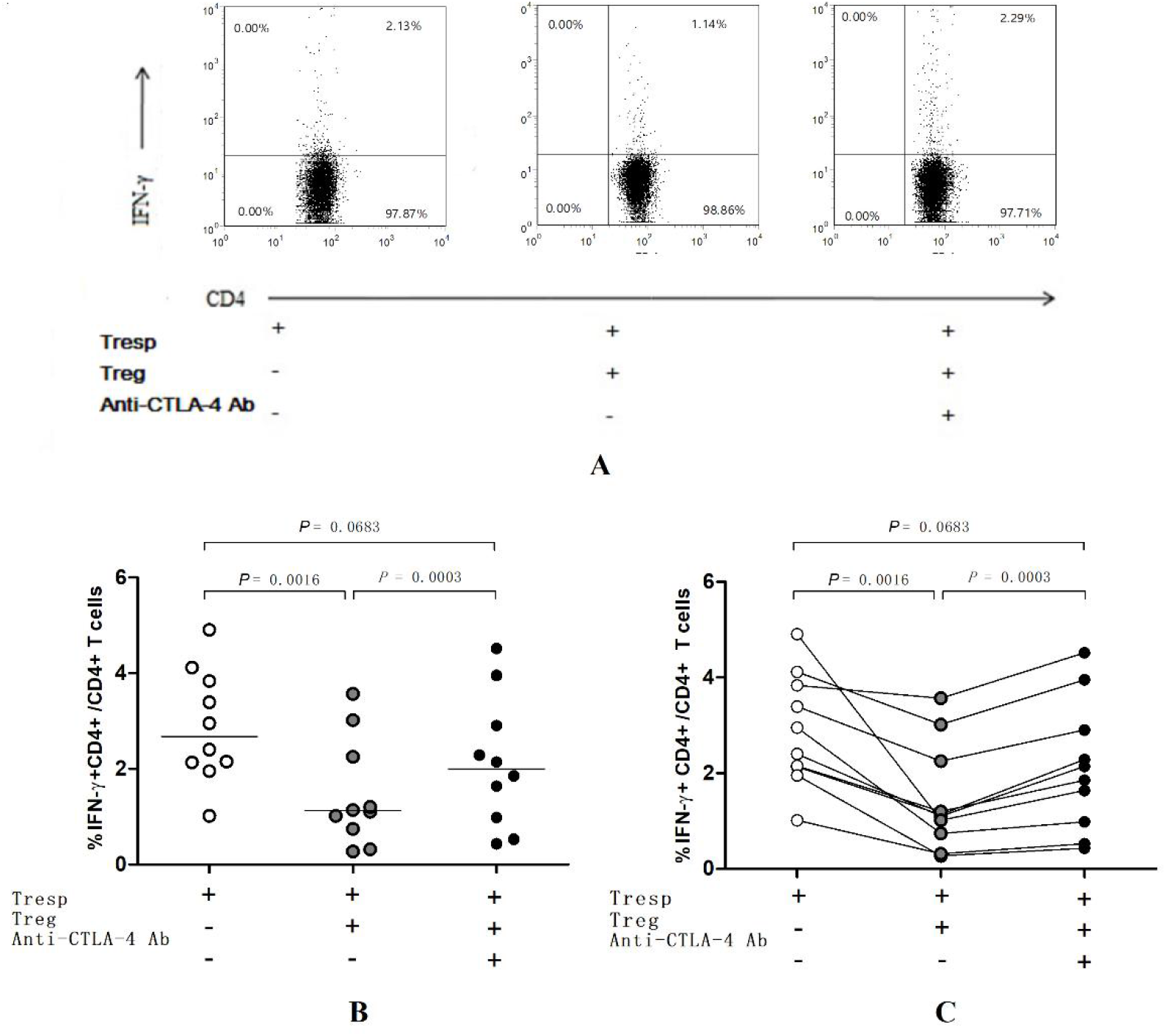
The CTLA-4 blockade abrogated the ability of Foxp3+ Tregs to suppress PPD-stimulated IFN-γ T effector response. (A) The representative flow cytometry histograms of IFN-γ-producing CD4+ T effector (Tresp, Foxp3-) cells under different culture conditions as indicated. Data gated on CD3 and CD4. (B) Frequencies of IFN-γ-producing Tresp with different conditions as indicated (n=10). The horizontal lines represent the medians for each culture condition. (C) The paired comparison of IFN-γ-producing Tresp under different culture conditions (n=10). Ten independent experiments were performed with comparable result. The horizontal lines represent the medians for each condition. No or very few IFN-γ+ CD4 T effector cells in medium control. The nonparametric quantitative variables across groups were compared by use of the paired t test. Significance was inferred for P<0.05 with two-sided.

We then examined whether the CTLA-4 blockade reversal of Treg inhibition in our PPD culture system indeed occurred as a result of decreased production of IL-10 by Treg cells. To our surprise, the CTLA-4 blockade did not lead to significant changes of IL-10 production by Foxp3+ Treg cells in the PPD culture system using paired comparison (*P*=0.1188, Figure S2).

Together, these results suggest that the CTLA-4 blockade abrogates the ability of Foxp3+ Treg cells to suppress TB PPD-stimulated IFN-γ T effector response, and such reversing effect is not correlated with any reduction of IL-10.

### Gene expression experiments demonstrated that the CTLA-4 blockade reversed the ability of Treg to suppress PPD-driven IFN-γ and IL-2 responses at mRNA levels, with no apparent changes in IL-10 and TGF-β

We then sought to examine if the CTLA-4 blockade abrogated the ability of Treg cells to suppress the PPD-stimulated cytokine responses at a transcriptional level. We employed real-time PCR assay to measure cytokine mRNA expression levels in the cells from the 3-day cultured system as described above. In the absence of CTLA-4 blockade, addition of Treg cells to the culture down-regulated IFN-γ and IL-2 expressions compared to the controls without Treg (*P*<0.0001 and *P*=0.0070, respectively). In contrast, the CTLA-4 blockade significantly reversed the ability of Treg to suppress IFN-γ and IL-2 expressions. The expression levels for these Th1 cytokines were significantly up-regulated in the presence of CTLA-4 blockade (*P*=0.0477 and 0.0459, respectively, Figure S3). However, the CTLA-4 blockage did not cause significant changes in inhibitory cytokines such as IL-10 and TGF-β under the culture conditions (Figure S3). These results provided the additional support suggesting that the CTLA-4 blockade abrogated the ability of Foxp3+ Treg cells to suppress TB antigen-driven Th1 cytokine responses.

### The CTLA-4 blockade significantly abrogated the Foxp3+ Treg suppression the proliferative response of PPD-specific T cells

Given the interrelation between Th1 cytokines and T-cell proliferation, we sought to examine if the CTLA-4 blockade also blocked the Treg suppression of PPD-driven T-cell proliferative response in our culture system. Thus, CD4+CD25-(Tresp) cells isolated from 5 active TB patients were labeled with CFSE, stimulated by PPD for 7 days in different Treg-Tresp ratio and assessed for CFSE-based proliferation in the absence and presence of CTLA-4 blockade. While addition of Foxp3+ Treg cells to the culture suppressed proliferation of PPD-specific Tresp cells, the CTLA-4 blockade reversed the Foxp3+ Treg suppression of PPD-driven proliferative T-cell response, reversing the proliferation rate from Treg-suppressive 28.9% up to 55.9% (*P*<0.05, Figure 5). Thus, these results suggest that CTLA-4 blockade could significantly abrogate the suppressive effect of Foxp3+ Treg cells on the proliferative response of PPD-specific T (Tresp) cells.

**Figure 5.**
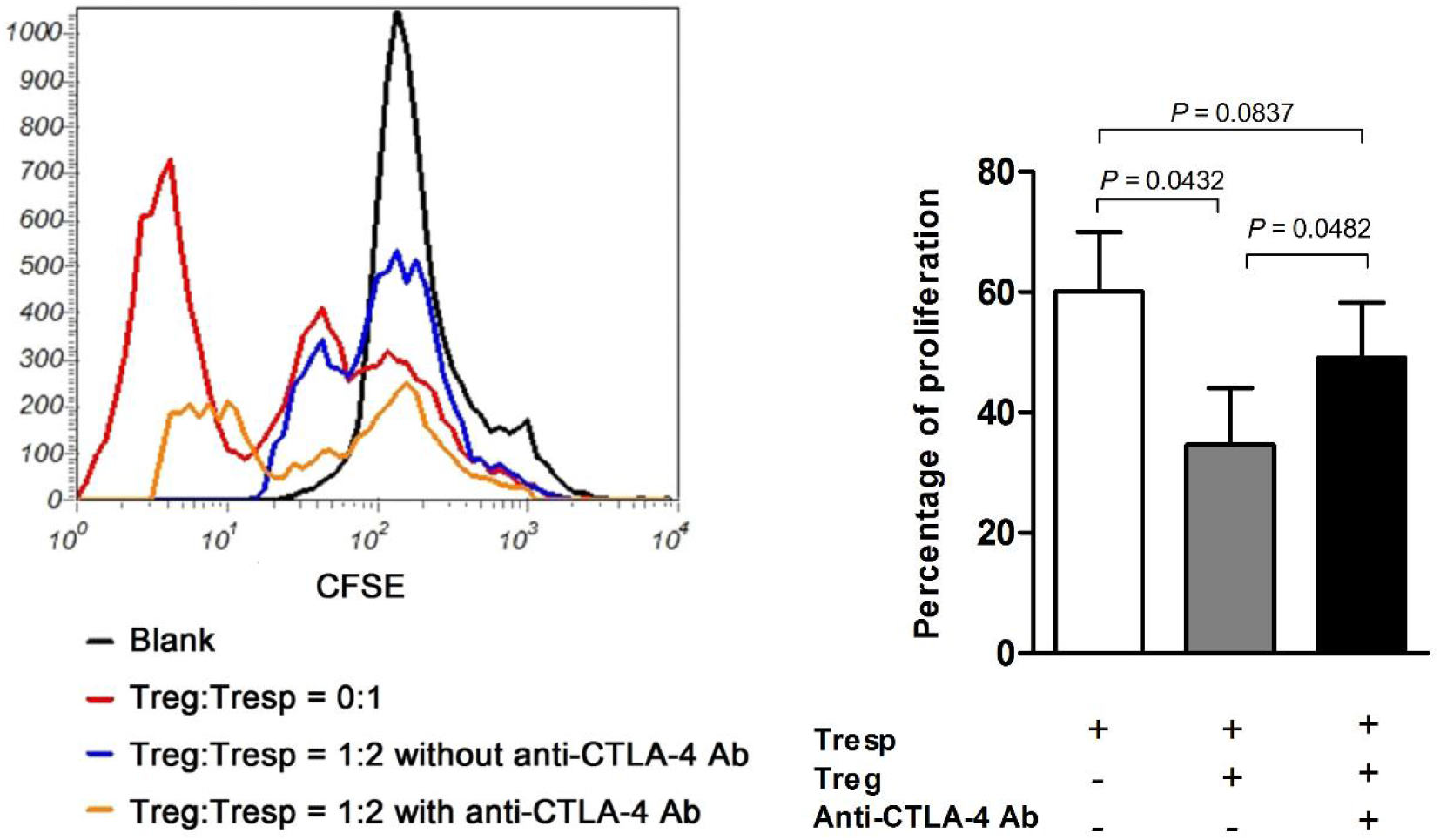
The CTLA-4 blockade significantly abrogated the suppressive effect of Foxp3+ Treg cells on the proliferative response of PPD-specific T cells. 5 active TB patients with positive sputum bacterium were enrolled for this assay. (A) The representative flow cytometry histograms of Tresp proliferation in response to 7-day PPD stimulation under different conditions. (B) Percentages of proliferating (divided) Tresp cells under different conditions (n=5). Five independent experiments were performed with comparable result. The nonparametric quantitative variables across groups were compared by use of the paired t test. Significance was inferred for P<0.05 with two-sided.

### The CTLA-4 blockade reversed the Treg suppression of the ability of T cells to restrict intracellular BCG and *M.tb* growth in vitro

Finally, we sought to determine if the CTLA-4 blockade could also reverse the Foxp3+ Treg suppression of the ability of T cells to restrict intracellular *M. bovis* BCG or *M.tb* H37Rv growth. To this end, Tresp cells and Foxp3+ Treg cells isolated from PBMCS of TB patients were added in different ratios to the co-culture containing BCG- or *M.tb*-infected macrophages in the presence or absence of the CTLA-4 blockade. Apparently, addition of Foxp3+ Treg cells to the culture had a suppressive effect on the T cell-mediated restriction of intracellular BCG and *M.tb* growth in autologous macrophages. Surprisingly, the CTLA-4 blockade reversed the Treg suppression of the ability of T cells to restrict intracellular BCG and *M.tb* growth, respectively (both *P*<0.01, Figure 6).

**Figure 6.**
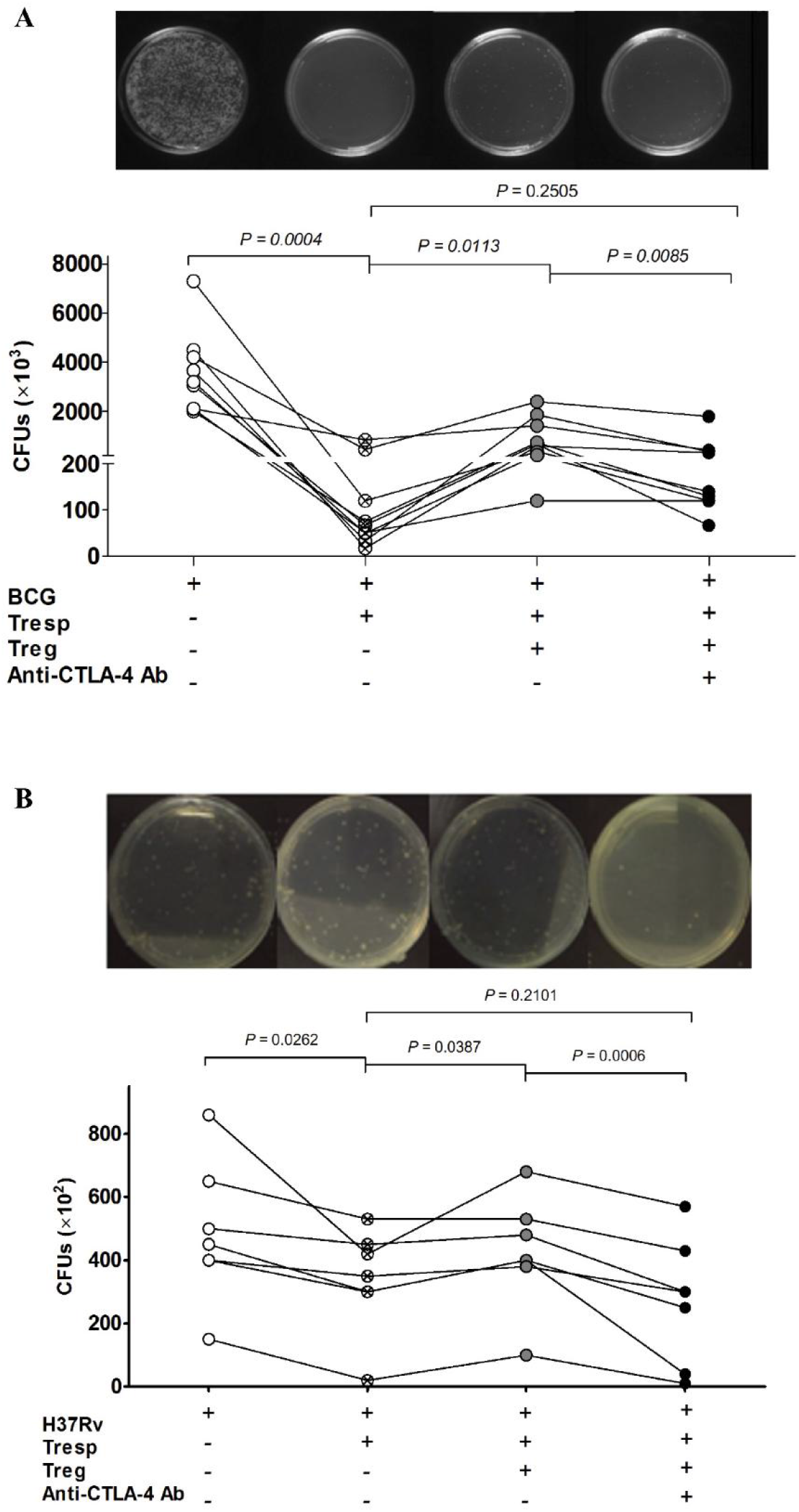
The CTLA-4 blockade reversed the Treg suppression of the ability of T cells to restrict intracellular BCG or *M.tb* growth in autologous monocyte-derived macrophages. (A) BCG CFU data from different co-culturing conditions of infected macrophages alone, Tresp and/or Foxp3+Treg in the presence or absence of the CTLA-4 blockade (cells purified from PBMCs of 9 TB patients, n=9). Nine independent experiments were performed with comparable result. (B) *M.tb* H37Rv CFU data from different co-culturing conditions of infected macrophages alone, Tresp and/or Foxp3+Treg in the presence or absence of the CTLA-4 blockade (cells purified from PBMCs of 7 TB patients, n=7).Seven independent experiments were performed with comparable result. BCG- or *M.tb* H37Rv-infected autologous macrophages were co-cultured with purified Tresp and Treg cells at the indicated ratio. At day 3 after the co-culture, cells were lysed and lysates were serially diluted and spread on 7H11 agar plates. CFUs were counted 2-3 weeks later. The nonparametric quantitative variables across groups were compared by use of the paired t test. Significance was inferred for P<0.05 with two-sided.

## Discussion

To our knowledge, the current study represents a first mechanistic investigation to address a potential role of CTLA-4 blockade in Foxp3+Treg suppression of TB antigen-specific T-cell responses in TB patients. The study allows us to make the following previously-unreported observations: (i). CTLA-4-expressing Foxp3+Treg cells are increased both in the circulation of pulmonary TB patients and in the pleural compartment of TB pleuritis; (ii). 6-month anti-TB treatment can significantly reduce TB-driven increases in CTLA-4+ Foxp3+ Treg subset; (iii). Antibody blocking of CTLA-4 pathway can reserve the ability of Foxp3+Treg cells to suppress anti-TB effector (IFN-γ) response of TB-specific CD4+ T cells at the cellular and transcriptional levels, and such reversing effects are not correlated with reduction of inhibitory cytokines IL-10 and TGF-β; (iv). The CTLA-4 blockade consistently antagonizes the Treg-mediated suppression of TB Ag-stimulated proliferative response; (v). Importantly, the CTLA-4 blockade also reverses the Treg suppression of the ability of T cells to restrict intracellular BCG and *M.tb* growth in macrophages. These new data may help to advance our understanding of the role of CTLA-4 blockade in Foxp3+ Treg suppression of anti-TB immune response or immunity.

Active human TB infection appears to significantly impact CTLA-4 homeostasis, leading to elevation of CTLA-4+Foxp3+ Treg cells. High frequencies of CTLA-4+Foxp3+ Treg cells are seen in the blood of active TB patients and also in the pleural fluid mononuclear cells and pleural membranes of patients with tuberculous pleurisy. Data implicate that CTLA-4+ Treg cells are increased in both systemic and local infection compartments during active TB infection. It is also noteworthy that anti-TB treatment can significantly restore imbalanced CTLA-4 hemostasis and reduce TB-driven high frequencies of CTLA-4+ Treg cells. Although increased frequencies of Foxp3+ Treg cells in TB have been reported(5, 30, 31), the current study shows new data of TB-driven increases in CTLA-4+Foxp3+ Treg subset. Data argue that CTLA-4 pathway may be involved in Treg suppressive function in TB.

Our findings in the mechanistic study suggest that CTLA-4 pathway is required for the ability of Foxp3+Treg cells to suppress TB antigen-specific T effector responses in active TB patients. This notion is supported by several pieces of experimental data in the current study. The CTLA-4 blockade apparently abrogates the capability of Foxp3+Treg cells to suppress PPD-specific IFNγ-producing CD4+ T effetcor responses. Similar blocking effects are also seen at the level of transcriptional responses of PPD-driven Th1 cytokines IFNγ and IL-2. Because Th1 cytokines are often involved in T-cell proliferation, it is not surprised that the CTLA-4 blockage can block the Treg-mediated suppression of proliferative responses of TB-specific T cells. Furthermore, the CTLA-4 blockage significantly reverses the Treg suppression of the ability of T cells to restrict or inhibit intracellular BCG and *M.tb* growth in macrophages. It has been shown in murine knock-out studies that specific CTLA-4 deficiency in Foxp3+ Treg cells impairs the Treg suppressive function in vivo and in vitro (13, 32, 33). Concurrently, CTLA-4 blockage has been shown to impact autoimmunity, inflammatory bowel disease, and tumor (34–36). However, whether CTLA-4 blockade can antagonize the Treg suppression of anti-TB immune responses in TB has not been demonstrated(37, 38). Our previous studies showed that TB-specific T cell effector subsets were depressed in TB patients (25, 26), and that mycobacterium-specific Th1 responses could be suppressed by cytokine-induced Foxp3+Treg cells (30). Now, the current study provides experimental data suggesting that CTLA-4 blockade can efficiently overcome Treg suppression of anti-TB T-cell immune responses.

Surprisingly, our results suggest that reversing effects of the CTLA-4 blockade on Foxp3+Treg suppression are not correlated with the reduced production of inhibitory cytokines IL-10 and TGF-β, while the role of IL-10 and TGF-β in the immunopathogenesis of tuberculous is still controversial(39, 40). Rather, our data suggest that reversing effects of the CTLA-4 blockade involve other mechanisms including cell-cell contact. Since the CTLA-4 blocking antibody used does not appear to directly enhance signaling for Th1 cytokine production (41), it is likely that antibody blocking of CTLA-4 interaction with its ligand B7 or other speculating complex molecules would efficiently reverse the Treg-mediated suppression of T-cell effector responses. Nevertheless, the current study does not have power to distinguish whether the reversing effects mainly result from CTLA-4 blockade on Treg cells, T effector cells or both. Elucidating the precise mechanism would require conditional knock-out platforms creating CTLA-4 deficiency selectively in primary human Foxp3+Treg or T effector cell subsets in TB patients. The sophisticated systems for manipulating selective primary human T-cell subsets are not easily available at this time, and such work may be beyond the scope of the current study.

Our data suggest that CTLA-3+Foxp3+ Treg cells and CTLA-4 pathway contribute to TB immunopathogenesis in humans. It is important to note that protective mechanisms or TB pathogenesis in humans remain largely unknown. Our findings in the CTLA-4 blockage experiments implicate that CTLA-4 pathway is required, at least in part, for the ability of Foxp3+Treg cells to suppress broad anti-TB T-cell responses at the levels of proliferation, Th1 effector functions, and intracellular *M.tb* growth control. Suppressions of these anti-TB T-cell responses presumably damage anti-TB immunity and ultimately lead to consequence of TB development or progression in humans. On the other hand, our data also support the view that the CTLA-4 pathway in TB is an important target for the immune manipulation. Since the CTLA-4 blockade in patients with selected infection or cancer has appeared promising (17, 19), rational or optimal CTLA-4 blocking antibody treatment may ultimately serve as potential adjunctive host-directed therapy against TB.

## Author Contributions

Lingyun Shao concepted and designed, analyzed and drafted the manuscript for important intellectual content; Yan Gao performed the main experiment and analyzed the data and drafted the manuscript; Xiaoyi Shao performed immunofluorescent staining and flow cytometric analysis; Shu Zhang performed the immunofluorescent staining, Qianqian Liu performed the in vitro intracellular *Mycobacterium* growth inhibition assay; Qin fang Ou, Bingyan Zhang, Qiaoling Ruan helped with collecting clinical datas and sample. Xinhua Weng and Wenhong Zhang designed and analyzed the study; Ling Shen and Zheng W. Chen designed, analyzed and drafted the manuscript for important intellectual content

## Correspondence

Wenhong Zhang: 12 Wulumuqizhong Road, Shanghai 200040, China.

